# Hippocampal neuronal synchronization in the rat - Dissociation of perception from associative learning

**DOI:** 10.1101/2022.12.20.521191

**Authors:** Kayeon Kim, Miriam S. Nokia, J. Matias Palva, Satu Palva

**Affiliations:** Helsinki Institute of Life Sciences, Neuroscience Center, University of Helsinki, Finland; Department of Psychology, University of Jyväskylä, PO Box 35, FI-40014, Finland; Department of Neuroscience and Biomedical engineering, Aalto University; Centre for Cognitive Neuroscience, Institute for Neuroscience and Psychology, University of Glasgow

**Keywords:** memory, classical conditioning, hippocampus, phase-locking, cross-frequency coupling

## Abstract

Learning requires flexible switching between perceiving and encoding external sensory stimuli, making associations with representations already stored in memory (retrieval) and acting accordingly to adapt to the environment. Hippocampus is known to be central in creating mnemonic representations, but how it could support the integration of these processes of memory encoding and retrieval during repetitive learning experiences has remained unknown. In this study, we recorded local-field potentials (LFPs) from the dorsal hippocampus during classical trace eyeblink conditioning in rats. An auditory conditioned stimulus (CS) was followed by a periorbital shock-unconditioned stimulus (US). From the LFPs, we separated hippocampal oscillations that were linked with the variation in the trial-by-trial processing of the CS from those linked with the gradual learning of the CS-US association over daily sessions of training. Short latency transient theta and gamma band oscillations were associated with learning so that both the oscillation amplitudes (strength) and phase locking increased as a function of training. In contrast, long-latency sustained alpha and beta oscillations were associated with perception of CS. These data show divergent oscillatory neural signatures for perception and learning in hippocampal subfields.

## Introduction

Episodic memory and spatial navigation are well known to be enabled by hippocampal– entorhinal (EC) circuits together with the neocortex (Buzsáki and Moser, 2013; Squire, 1992). Memory formation and synaptic plasticity has been related to hippocampal theta (θ, 3–12 Hz) oscillations (for reviews see Buzsáki, 2002; Colgin, 2013, 2016). The presence of dominant θ oscillations in the hippocampus prior to training predicts more efficient learning in a subsequent task in rats (Nokia et al., 2012), rabbits (Berry and Thompson, 1978; Nokia et al., 2009; Seager et al., 2002) and in humans (Caplan et al., 2003). During associative learning in rodents, EC, and hippocampus and striatum are coupled both via θ (DeCoteau et al., 2007; Takehara-Nishiuchi et al., 2012) and beta (β) / gamma (γ) band (20–40 Hz) phase-synchronization (Colgin et al., 2009; Igarashi et al., 2014). It also showed relation with cross-frequency coupling (CFC) between θ phase and β / γ oscillations amplitudes across striatum and hippocampus (Tort et al., 2008) and it’s evident that both in CA3-CA1 and EC-CA1 circuits involve CFC between θ phase and γ amplitude (Fernández-Ruiz et al., 2017). Further, θ oscillations synchronize during eyeblink classical conditioning in the hippocampus and the cerebellum, the structure responsible for the execution of the (learned) behavioral responses (Wikgren et al., 2010).

Adaptive behavior change requires multiple processes from perception and encoding of external stimuli to associating this with representations of earlier experiences and finally to the selection and execution of an appropriate motor response. It is thought that the role of the hippocampus in this process is to form an initial snap-shot representation of the multimodal information of an external event (Buzsáki, 1989) and to link disparate neocortical representations into a coherent memory trace (Lisman, 1999; Wallenstein et al., 1998) based on an assumed hippocampal memory index (Teyler and Rudy, 2007). Clinical work shows that formation of new episodic memories is impaired after hippocampal damage (Scoville and Milner, 1957) and that memory-impaired patients with medial temporal lobe lesions also exhibit diminished markers for conscious perception (Urgolites et al., 2018) as well as show attenuated and temporally more dispersed responses to visual stimuli (Reber et al., 2017). This suggests that hippocampal circuits may play a central role not only in memory formation but already in perception (Kreiman et al., 2002). In fact, recent work utilizing human intracranial and scalp electroencephalogram (EEG) suggested that the hippocampus could represent both sensory and mnemonic information and act as a switchboard between internal and external representations (Treder et al., 2021).

In this study, we set out to investigate whether electrophysiological oscillations in the hippocampus could dissociate between the on-the-spot processing of external stimulus and adaptive behavior based on it (hereafter, perception) and the gradual acquisition of an association between stimuli leading to adaptive behavior change over time (hereafter, associative learning). To this end, we recorded local-field potentials (LFPs) from the dorsal hippocampus in freely-moving adult healthy Sprague-Dawley male rats during classical trace eyeblink conditioning (Clark et al., 2002). The task requires representation of the tone-conditioned stimulus (CS), learning of an association between the CS and the unconditioned stimulus (US, here a shock to the eyelid) and finally an adaptively timed conditioned response (CR), namely an eyeblink in response to the CS. In humans, correct stimulus-response (CS-CR) mapping and conscious perception of the association between the CS and the US are thought to be a prerequisite for successful learning (Clark and Squire, 1998). Here we used the CR as an index of perception (see above and Methods for a definition) and a change in the proportion of CRs, the hit rate (HR), across sessions as a measure of learning.

## Results

### LFPs were recorded from the hippocampus during trace eyeblink conditioning in rats

We recorded LFPs (1–500 Hz) from the hippocampus (Fig. 1A) of adult male Sprague-Dawley rats (n = 8) during trace eyeblink classical conditioning task (Fig. 1B, see Methods for more details). We used self-made wire-electrodes with large contact surface (> 50 microns) and tips separated by 200–250 microns, and recorded LFPs reflecting joint activity of thousands of neurons from both fissure (f) and from the hilus (h). Fissure encompasses the hippocampal region occupied by the dendrites of the dentate gyrus (DG) granule cells receiving main input from the entorhinal cortex (EC), mammillary bodies and the brainstem. Hilus defines the regions occupied by passing axons form DG granule cells conveying the signals to CA3 pyramidal cells (Fig. 1A) (Scharfman, 2016).

**Figure 1.**
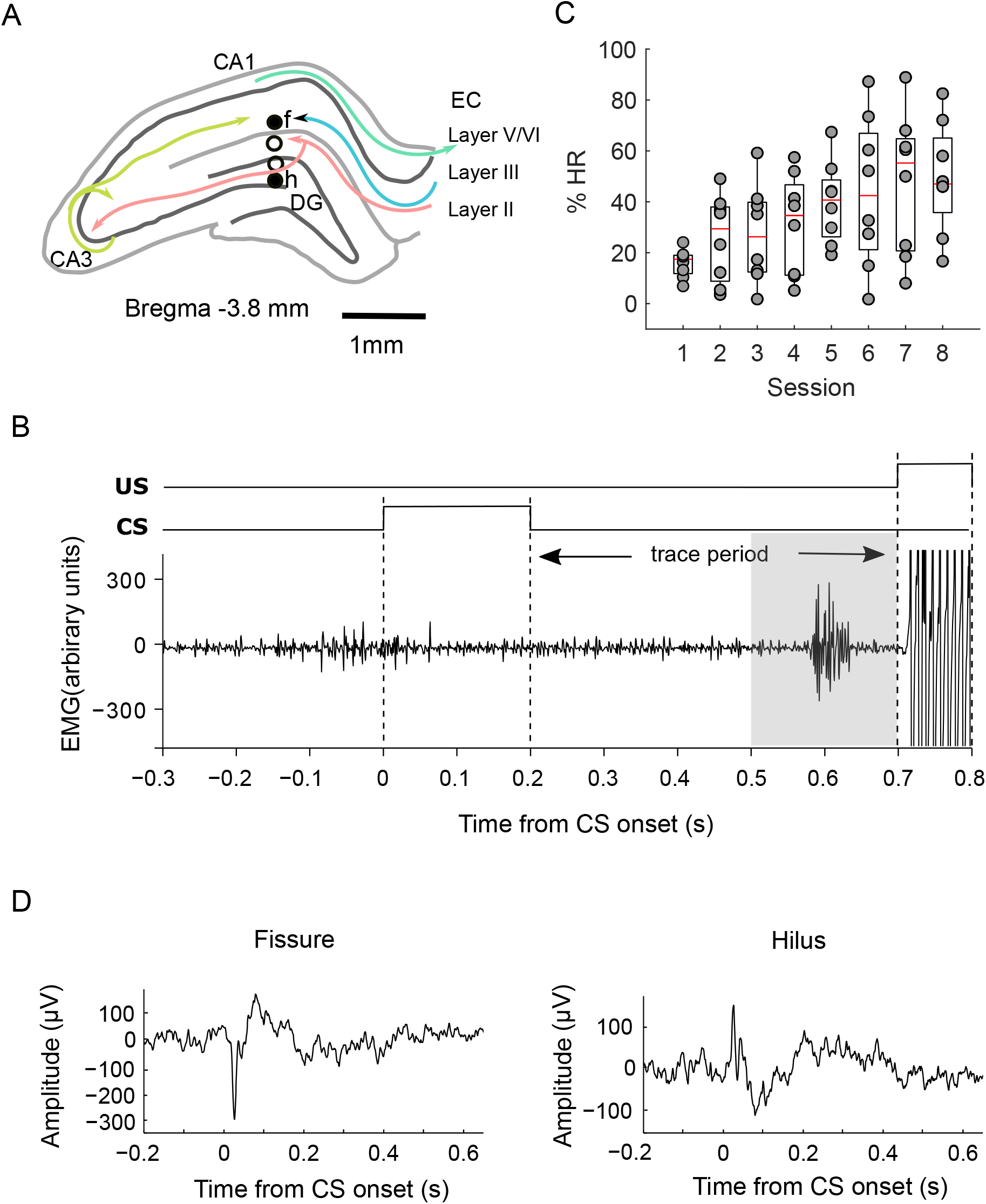
Local-field potentials from the hippocampus during trace eyeblink conditioning in adult male Sprague-Dawley rats. **A**. Graphical representation of recording electrode placement in the rat hippocampus. Target sites, fissure (f, upper) and hilus (h, lower) are shown. **B**. An example of electromyogram (EMG) trace from the eyelid during the task, trace eyeblink conditioning. The gray area indicates the time window when eye blinks were accounted as a conditioned response. A tone was used as conditioned stimulus (CS) and a periorbital shock was used as the unconditioned stimulus (US). **C**. Behavioral performance (%HR) plotted against session. Individual rats (n = 8, circles) on top of box plot. **D**. A representative peri-stimulus LFP trace aligned to CS onset, averaged across all trials (n = 60) from one example session of one rat in each recording site; fissure (left), hilus (right).

Eight trace eyeblink conditioning sessions, one per day, were conducted for each rat. During the daily session, 60 paired (CS + US) trials were presented. Each trial started with a 200-ms tone-CS and was followed by a 500-ms stimulus-free trace period, and then a 100-ms shock-US to the eyelid. Eyeblinks were recorded using bipolar wire-electrodes implanted in the eyelid (electromyogram, EMG) (Nokia et al., 2017).

### Rats learned trace eyeblink conditioning

The last 200 ms of the trace period was used to assess whether a CR (eye blink prior to US onset) occurred in each trial (see Fig. 1B for an example). First, we approximated the efficacy of CS perception based on the CRs and defined a hit rate indicating the percentage of trials during which a correctly timed blink (a CR) was performed. Then, we defined learning as an increase in HR as the conditioning progressed from session one to eight. The percentage of CRs significantly increased as a function of session (Pearson r = 0.49, *p* = 4.51×10^−5^) (Fig. 1C), and a significant difference in HR among sessions (repeated measures ANOVA: *F*_[7, 49]_ = 5.72, *p* = 7.17×10^−5^) indicates that the rats learned to anticipate the shock-US as training progressed.

### Spatio-temporal dynamics of oscillations in response to CS

Next, we characterized hippocampal oscillatory activity in response to the CS. The evoked LFP showed a transient peak in both fissure and hilus (Fig. 1D). Time frequency representation (TFR) of oscillation amplitudes showed a transient wide-band increase in oscillation amplitudes at 5–120 Hz in response to the CS in both hilus and fissure (Fig. 2A). The transient response was followed by sustained oscillations in the 8−12 Hz range, here termed alpha (α) band for consistency with human literature, accompanied with a sustained response in high-gamma (high-γ, 50–100 Hz) band and a concurrent suppression of oscillation amplitudes in the θ and in the β to low-γ -bands. Oscillation amplitudes averaged across three time windows (See Methods), showed that only the oscillation amplitudes transiently increased up to 480 Hz in both fissure and hilus were significant compared to baseline (Fig. 2B, red trace, Monte-Carlo *p* < 0.025, two-tailed, False Discovery Rate (FDR) corrected). The sustained α and γ band oscillations were significantly increased compared to baseline only in fissure but not in hilus, while β oscillations were suppressed both in fissure and hilus (Fig. 2B green, blue trace, Monte Carlo *p* < 0.025, two-tailed, FDR corrected). Despite similar spatiotemporal oscillatory profiles in hilus and fissure, sustained oscillatory response amplitudes to CS were stronger in fissure (Fig. 2B, green trace, Monte-Carlo *p* < 0.025, two-tailed, FDR corrected) in accordance with fissure being the primary site of EC inputs via the perforant pathway (Scharfman, 2016). Next, we characterized whether oscillations would also show phase locking of ongoing oscillations to stimulus onset (inter-trial phase-locking) estimated with phase-locking factor (PLF) (Palva et al., 2005). Like oscillation amplitudes, transient wide-band PLF characterized LFP in both fissure and hilus (Fig. 2C) this being significant (Fig. 2D, Monte-Carlo *p* < 0.025, two-tailed, FDR corrected) and indicating that oscillations were robustly phase-locked to CS onset.

**Figure 2.**
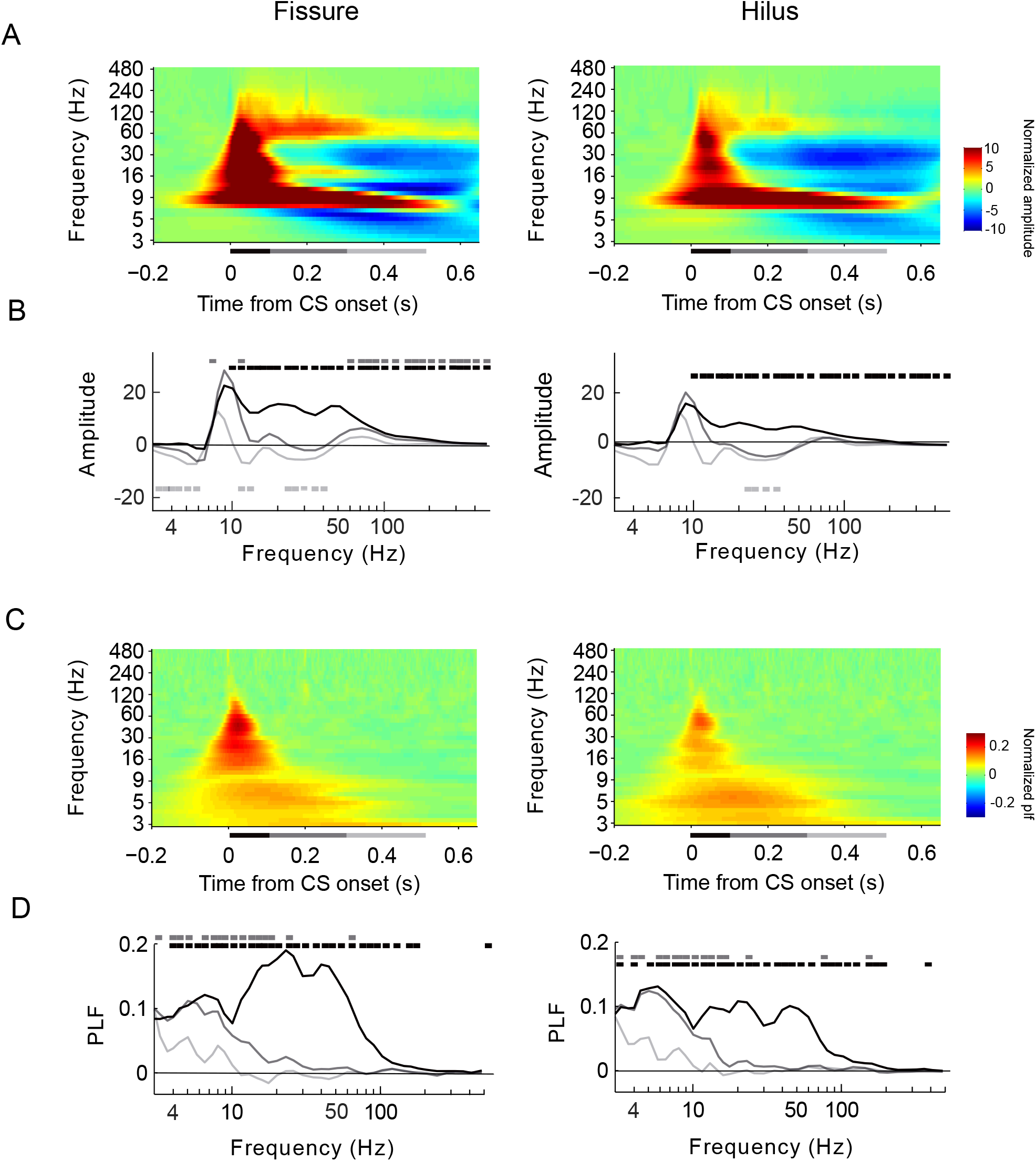
Spatio-temporal dynamics of hippocampal oscillations during trace eyeblink conditioning. **A**. Baseline-corrected time-frequency representation (TFR) of oscillation amplitudes (A) in response to CS separately for fissure (left) and hilus (right). Oscillation amplitudes were averaged across trials, sessions, and then across all rats (n = 8). The horizontal bar below the panel indicates analysis periods relative to CS onset; 0–0.1 s (black), 0.1–0.3 s (dark gray), 0.3–0.5 s (light gray). **B**. Oscillation amplitudes averaged across three time windows. Horizontal colored bars denote statistical significance at *p* < 0.025 (see Materials and Methods for details). **C**. Phase-locking factor (PLF) as in A. **D**. PLF as in B. Horizontal bars denote statistical significance at *p* < 0.025.

### Sustained increase of α and β oscillation amplitudes is associated with CS perception

Here we approximate the eyeblink-CR to reflect accurate perception of the CS, retrieval of the previously acquired representation of the US and the motor response that follows it, namely the eyeblink. For simplicity, we refer to this whole process as perception. To assess whether distinct spatiotemporal patterns of oscillations would be correlated with perception, we computed oscillation amplitudes and PLF separately for trials with a CR and those without a CR and further computed the difference in oscillation amplitudes (Fig. 3A) and PLF (Fig. 3B) between the CR and the no-CR trials. We found no effect of perception on transient oscillation amplitudes. However, modulation of sustained oscillation amplitudes in α, β and γ bands was stronger for the CR than for no-CR trials in both fissure and hilus, (Fig. 3A, Monte Carlo *p* < 0.050) so that these were stronger for the perceived CR. In contrast, there was no noticeable difference between perceived and unperceived trials in the PLF (Fig. 3B).

**Figure 3.**
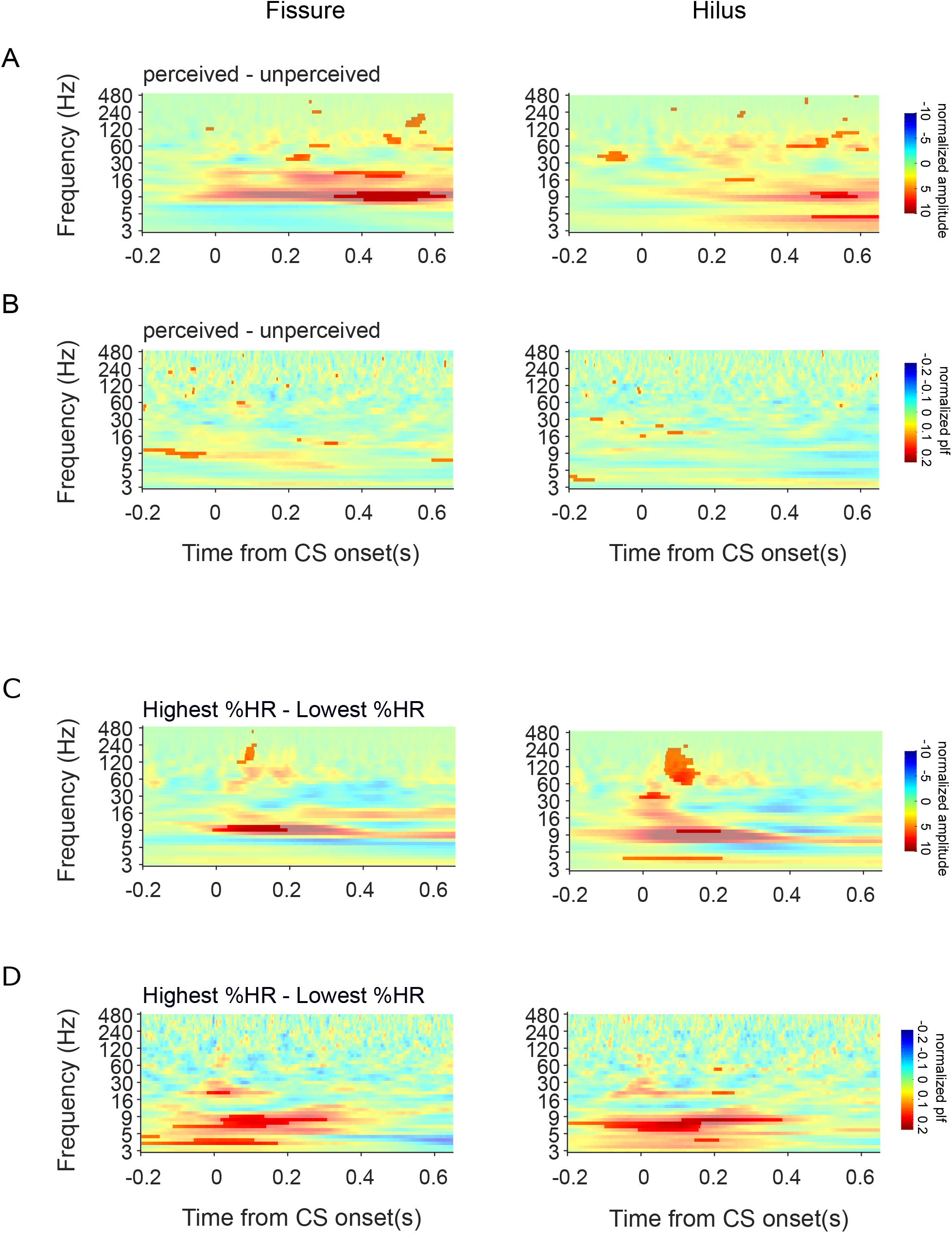
Perception and learning are associated with distinct spatio-temporal profiles of hippocampal oscillations. **A, B**. A) Oscillation amplitudes and B) PLF between perceived and unperceived trials, left (fissure) and right (hilus). Significant differences between perceived and unperceived (Monte-Carlo *p*<0.05) are masked. **C. D**, C) Oscillation amplitudes and D) PLF for the difference between highest %HR and lowest %HR.

### Transient early oscillatory activity predicts associative learning

We then tested whether we could dissociate a separate spatiotemporal pattern for the learning across the sessions by analyzing the data from the sessions with the highest HR and the lowest HR (See Methods for more details) and for their difference (Fig 3C, amplitude; 3D, PLF). Transient α and γ oscillation amplitudes in both fissure and hilus, and θ band amplitude in hilus during first 200– 300 ms after CS onset (Fig. 3C, Monte Carlo *p* < 0.050) predicted learning. In contrast to perception, also α -band PLF in both fissure and hilus, and θ-band PLF in fissure also predicted learning (Fig. 3D, Monte Carlo *p* < 0.025). Interestingly, learning was also predicted by pre-stimulus θ−band PLF in fissure and α−band PLF in hilus (Fig. 3D, Monte-Carlo *p* < 0.025) suggestive of the presence of anticipatory phase-reorganization for the sessions with learning stimulus associations.

### Intra-hippocampal phase synchronization and directional interactions predicts perception and learning at different latencies

Oscillations in θ and γ bands are widely observed in hippocampal sub-regions (Belluscio et al., 2012; Rangel et al., 2016). In the hippocampus, information flows from EC to dendrites of the DG granule cells and CA1 pyramidal cells lining the fissure, and from the DG to CA3 pyramidal cells via the hilus. From hilus information feeds back to the CA1 pyramidal cell dendrites lining the fissure (see Fig. 1A). Taken that oscillatory synchronization is thought be a mechanism for regulating communication (Fries, 2015) we studied whether oscillations between hilus and fissure would show phase-synchronization. Robust phase synchrony between hilus and fissure in the α, and high-γ frequency bands was sustained after CR (Fig. 4A, right, Monte-Carlo *p* < 0.010, FDR corrected) despite the lack concurrent oscillation amplitude increases. However, we found no correlation of synchronization with perception or with learning (i.e., the increase in HR over sessions) (results not shown).

**Figure 4.**
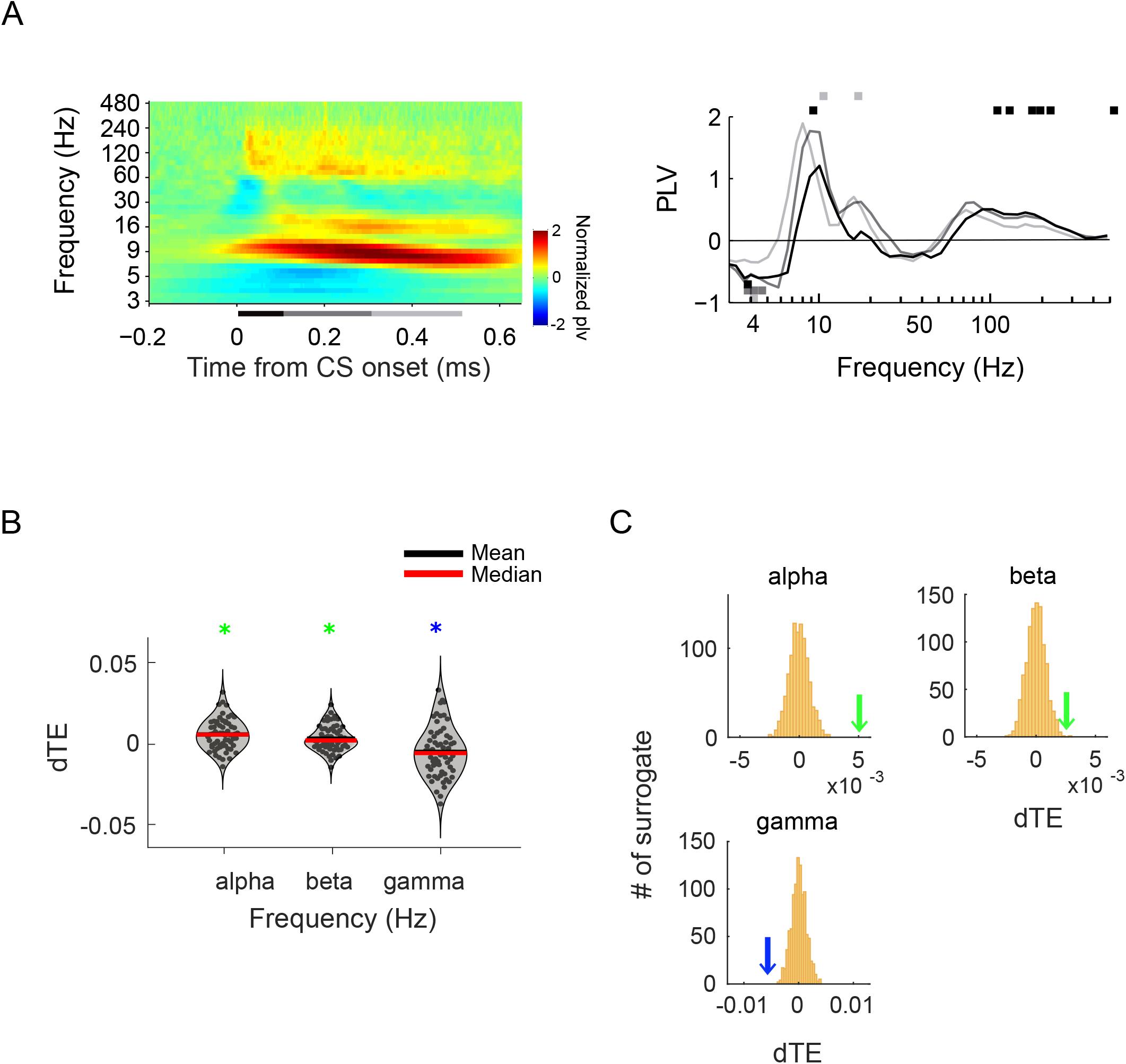
Inter-areal synchronization and directed interactions between hilus and fissure. **A**. Inter-areal synchronization as estimated with phase-locking value (PLV) averaged across trials, sessions, and then across all rats (left). Horizontal bars denote significance level *p*< 0.025 (right). **B**. Phase transfer (dTE) across sessions and rats. Green marks indicate significant dTE from hilus to fissure, blue mark indicates significant dTE from fissure to hilus. **C**. Distribution of mean dTE derived from surrogate data by trial shuffling. The green or blue arrows in each figure indicate mean dTE of the empirical data.

To establish whether oscillatory interactions would be associated with feed-forward information flow from fissure to hilus or feed-back processes from hilus to fissure, we next estimated the directionality of synchronization using phase transfer entropy (pTE) (Lobier et al., 2014) focusing on the frequencies showing significant inter-areal synchrony (See Fig. 4A). Differential TE (dTE) was derived by difference between pTE_hil→fis_ and pTE_fis→hil_ across all sessions and rats. In α and β bands, information flow was significant from hilus to fissure (Fig. 4B-C green marks, Monte-Carlo *p* < 0.050) whereas oscillations at high-γ (80 Hz) band showed the opposite direction, from fissure to hilus (Fig. 4B-C, blue marks, Monte-Carlo *p* < 0.05).

### Nested theta-gamma oscillation associated with perception and learning

As theta oscillations are thought to underlie a central clocking mechanism for timing of gamma oscillations, we further mapped the phase amplitude coupling (PAC) or “nesting” of these oscillations (Canolty and Knight, 2010; Colgin, 2011; Fell and Axmacher, 2011; Hyafil et al., 2015; Lisman and Jensen, 2013; Schroeder and Lakatos, 2009) across frequencies and laminar pairs. Robust θ/α to γ PAC characterized LFP signal within hilus and between hilus and fissure across all analysis time windows tested (Fig. 5A-C, t-test against zero, *p* < 0.050, FDR corrected), confirming the presence of the theta-gamma code in classical eyeblink conditioning task. In addition, transient PAC (analysis window 0–100 ms after the CS onset) between low-γ (30–45 Hz within hilus, 35–45 Hz between hilus and fissure) to higher γ (>180 Hz) frequencies was found in hilus and between hilus and fissure (Fig. 5A, t-test against zero, *p* < 0.010, FDR corrected). Even though we did not find significant link between 1:1 inter-areal synchrony with behavioral performance, critically, the strength of θ/α–γPAC during a period >300 ms from CS onset was linked with both perception and learning (Figure 5A-C, correspondingly, paired t-test, *p* < 0.01), the association between strength of PAC being prominent during earlier latencies than that for perception.

**Figure 5.**
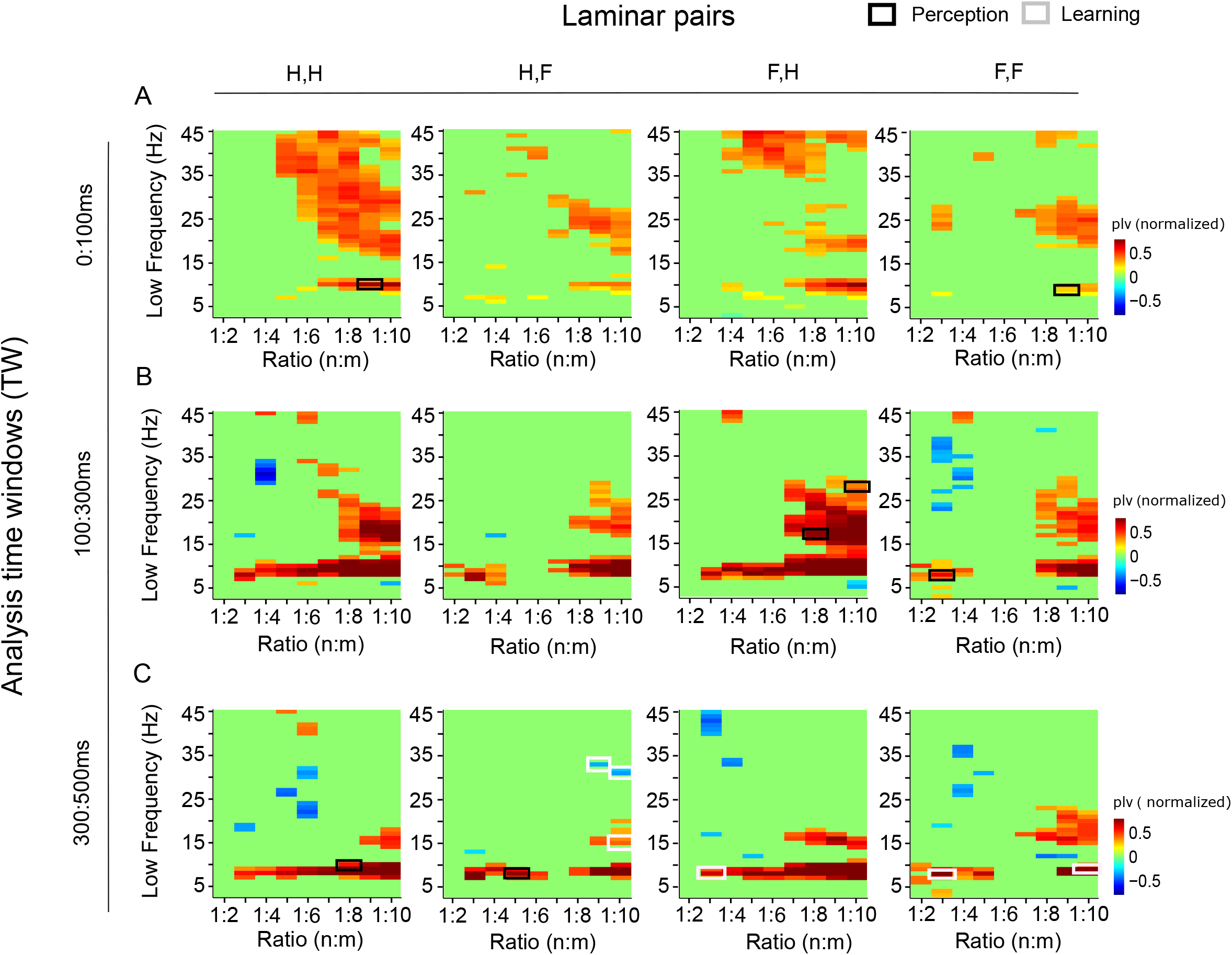
Phase amplitude coupling (PAC) for CS. PAC is plotted for each laminar pair (H-hilus; F-fissure) and for each analysis window of interest (rows from top to bottom **A-C**; A, 0-0.1s; B, 0.1-0.3s; C, 0.3-0.5s after stimulus onset). Y-axis shows the lower frequency (Hz) used for computing *n:m* PAC and is plotted against the corresponding *m:n* ratio. Each panel shows significant PLV difference between post-stimulus and pre-stimulus time-windows (*p*<0.05, FDR corrected) where the black rectangles indicate significant difference between perceived and unperceived and white between highest %CR and lowest %CR (paired t-test, *p*<0.025).

## Discussion

Trace eyeblink conditioning has been used widely to study neuronal mechanisms of learning in humans (Cason, 1922), rabbits (Schneiderman et al., 1962), and in rodents (McEchron and Disterhoft, 1999; Tseng et al., 2004; Weiss et al., 1996) and have resolved a fundamental functional role of hippocampal θ oscillations in associative learning. However, the task also requires conscious awareness of the stimulus contingencies between CS and US during the trace period that allow the learning the of stimulus association (Cheng et al., 2008; Clark and Squire, 1998). Despite this critical role of sensory perception and memory retrieval in the trace eyeblink conditioning task, how hippocampal circuits represent conscious perception of the CS and its correct mapping with US-eyeblink stimulus-response mapping, have remained unknown.

In this study, we used a classical trace eyeblink conditioning task to dissociate the hippocampal oscillation dynamics associated with perception (and correct stimulus-response mapping) from those dynamics associated with learning i.e., improvement in performance across sessions. In line with the hypothesis, our results demonstrated a presence of distinct hippocampal oscillation dynamics is associated with perception and associative learning.

The auditory-CS was followed by a transient wide-band increase in hippocampal oscillation amplitudes both in fissure and hilus. This response was similar to that found in the sensory cortices in humans (Hirvonen and Palva, 2016; Julku et al., 2021; Palva et al., 2005, 2011) and primary auditory cortex in guinea pigs (Voigt et al., 2018). The strength of this transient response was not correlated with perception but, nevertheless, the strength of θ/α and γ band oscillation amplitudes predicted learning across sessions so that better learners showed stronger responses. In contrast, the subsequent sustained α, β and γ band responses in fissure and hilus were correlated with stimulus perception, but not with learning across daily training sessions. These results demonstrate that separate hippocampal oscillation dynamics are associated with perception and associative learning, similar to that observed with electrophysiological recordings during a visual associative memory recall task in humans (Treder et al., 2021). Our results show that the sequential pattern of hippocampal oscillation dynamics could be a mechanism for mediating learning of stimulus associations in classical conditioning.

### Transient early oscillations and synchronization predict learning but not conscious perception

The correlation of transient oscillation amplitudes with learning across daily sessions observed in our current study is in line with the traditional view of θ amplitude (Berry and Thompson, 1978; Winson, 1978) and θ phase (Nokia et al., 2015) correlating with learning. Furthermore the results show that the strength of neural activity around stimulus onset rather than at later latencies predicts subsequent memory formation in Pavlovian conditioning task in rodents similar to that found for single- and multi-unit activity recorded from the human hippocampus during recognition (Urgolites et al., 2020) and episodic (Griffiths et al., 2021) memory tasks. Also, phase-synchronization within the hippocampus showed temporal dissociation for learning and perception as the early γ-band synchronization predicted learning but not perception. The lack of correlations with CS perception suggest that the early transient CS-induced oscillations and synchronization may reflect unconscious feed-forward processing (Lamme and Roelfsema, 2000), possibly relaying the information for intrahippocampal processing, and demonstrate that learning in this Pavlovian conditioning task is partially dependent on the strength of fast automatic unconscious processing in the hippocampus.

### Stimulus perception and response mapping are associated with strengthened oscillations

Beyond encoding of physical environment such as spatial mapping during navigation or exploration (Morgan et al., 2011; O’Keefe and Dostrovsky, 1971; O’keefe and Nadel, 1978) hippocampus is also related to representing and encoding of visual information in primates (Jutras et al., 2013; Lee et al., 2012; Zeidman et al., 2015), and odors in rodents (Kay, 2014). Here, sustained α and γ band responses in fissure and hilus as well as suppression of β band amplitudes in hilus were correlated with stimulus perception, hence supporting the role of the medial temporal lobe structures in conscious perceptual operation (Cheng et al., 2008). These oscillations were, however, not associated with learning and training effects across daily training sessions indicating that perceptual processes are distinct from those underlying learning and occurring at later latencies when perceptual information is broadcasted in a widespread brain network (Dehaene and Changeux, 2011; Del Cul et al., 2007; Lamy et al., 2009).

Previous studies have linked hippocampal oscillations to encoding stimulus associations in rodents (Zheng et al., 2016), monkeys (Jutras et al., 2009) and humans (Griffiths et al., 2021, 2019), but not to encoding perceptual representations *per se*. Our results, however, suggest that hippocampal oscillation dynamics support neocortical oscillations not only in learning but also in the perceptual processes. Interestingly, perceptual modulations in oscillation amplitudes detected in our current study correspond to latencies where stimulus-evoked responses are correlated with conscious awareness in human EEG data (Del Cul et al., 2007; Sergent et al., 2005). The timing was also similar to that of perception from external stimulus being converted to internal representation observed in human mnemonic association task (Treder et al., 2021). Taken together, we propose that the switch from transient wide-band oscillations to sustained θ, α and γ synchronization likely reflects a switch from the unconscious learning-induced feed-forward processing to perceptually relevant processes.

### Synchronization between hippocampal subregions and CFCs across frequencies

Local oscillations were also accompanied by perceptually relevant θ−α, β and γ band synchronization within the hippocampus in line with the framework that oscillatory synchronization is a mechanism for regulation of neuronal communication across spatially distributed assemblies (Fries, 2015). These results are well in accordance with well-established anatomical connectivity between hippocampal subregions (Andersen et al., 2006) that show also frequency-dependency (Akam et al., 2012; Das and Menon, 2021; Gloveli et al., 1997). For high-α, β, and low-γ oscillations the direction of information flow was significant from hilus to fissure while the opposite was observed for high-γ activity. Taken that neocortex sends projections to DG granule cells whose dendrites occupy the molecular layer along the fissure, and that the DG granule cells convey the signal to CA3 pyramidal cells via their axons passing in the hilus, these results suggest that in accordance with previous work (Jensen et al., 2015; Michalareas et al., 2016; Richter et al., 2017) synchronization in γ−band could mediate feed-forward processing of information while the slower frequencies could exert the top-down control for the memory-based formation of associations, mediating feed-back from EC to DG granule cell and CA1 pyramidal cell dendrites.

In accordance with θ oscillations yielding a clocking mechanisms for temporal coding of information in hippocampo–subicular network (Buzsáki and Chrobak, 1995; Chrobak and Buzsáki, 1996; Lisman and Jensen, 2013; Wang and Buzsáki, 1996; Whittington et al., 1995), robust PAC between phase of θ-α oscillation and amplitude envelope of γ oscillation characterized local and inter-areal hippocampal oscillation dynamics. These data are in line with prior studies that have observed θ-γ PAC in rat hippocampus during a T-maze task (Tort et al., 2008), associative learning (Tort et al., 2009) and during exploration and rapid-eye movement sleep (Belluscio et al., 2012; Scheffer-Teixeira et al., 2012; Tort et al., 2010) and in neocortex during a recognition memory task (Bahramisharif et al., 2018). Importantly, the strength of PAC between θ−α to γ oscillations predicted learning across sessions during the early latencies and predicted perception during the later latencies similarly to local oscillation amplitudes and PLF. This speaks for the importance of temporal precision of stimulus processing on learning perhaps via spike-time -dependent plasticity (Fell and Axmacher, 2011; Mansvelder et al., 2019; Singer, 1995; Uhlhaas et al., 2010) that allows forming of the stimulus associations.

## Conclusions

We used a classical trace eyeblink conditioning task to dissociate the hippocampal oscillation dynamics associated with perception and correct stimulus-response mapping from those associated with learning. We show distinct spatiotemporal patterns of oscillations for these functions. Perceptual processes were associated with late oscillation amplitude modulations and synchronization while associative learning was correlated with early transient responses. These results demonstrate the presence of distinct hippocampal oscillation dynamics associated with perception and associative learning.

## Materials and Methods

### Ethical statement

The experiment was conducted in agreement with directive 2010/63/EU of the European Parliament and of the Council on the care and use of animals for research purposes. The experiments were approved by the Regional State Administrative Agency of Southern Finland. The ARRIVE guidelines (https://arriveguidelines.org/arrive-guidelines) were followed.

### Subjects

Adult healthy male Sprague-Dawley rats were used as subjects. We analyzed data from eight rats, which belonged to a normal control group out of three different experimental groups that were used to study the effects of hippocampal stimulation on associative learning effect while rats were performing trace eyeblink classical conditioning task (Nokia et al., 2017).

### Experimental procedures

#### Surgery

Surgical procedures are described in detail in Nokia et al., (2017). Briefly, rats were anesthetized with i.p. injection of pentobarbital (60 mg/kg) and treated for pain with carprofen (5 mg/kg, s.c.) and buprenorphine (0.03 mg/kg, s.c.). Two bundles of 4 wire (Formwar insulated nichrome, bare diameter 50 microns) electrodes glued together with a tip separation of 200 to 250 microns were implanted into the dorsal hippocampus aiming at the dentate gyrus (3.6–4.5 mm posterior, 1.5–2.2 mm lateral and 3.6–4.0 mm below bregma) (see Figure 1A). Skull crews served as reference (11 mm posterior and 2 mm lateral to bregma) and ground (4 mm anterior and 2 mm lateral to bregma). Note that bipolar wire stimulation electrodes were lowered to the ventral hippocampal commissure (1.3 mm posterior, 1.1 mm lateral to bregma and 4.1 mm below bregma) but were not used in the animals included in this study. To stimulate the eyelid and record electromyography (EMG) during conditioning, two bipolar electrodes made of stainless-steel wire insulated with Teflon (bare diameter 127 microns) were implanted through the upper right eyelid. At last, the whole construction was secured in place using dental acrylic cement. Each rat was allowed to recover for at least one week and medicated for pain with buprenorphine.

#### Recordings

The EMG signal was band-pass filtered between 100 and 300 Hz. To acquire LFPs, a low-noise pre-amplifier (10 x) was directly attached to the electrode connector in the rat’s head. The LFP signals were filtered (1–5000 Hz) and further amplified (50 x), then digitized at 20 kHz and low-pass filtered at 500 Hz. Finally, all signals were stored at a 2-kHz sampling rate (USB-ME-64, Multi-Channel Systems). For details, see Nokia et al. (2017).

#### Histology

Rats were euthanized by exposure to a rising concentration of CO_2_ and then decapitated. The locations of the electrode tips in the brain were marked by passing a DC anodal current (200 mA, 5 s) through them. The brain was then removed, fixed in 4% paraformaldehyde solution and coronally sectioned with a vibratome (Leica VT1000). The slices were stained with Prussian blue and cresyl violet. The electrode tip locations were determined with the help of a conventional light microscope and a brain atlas (Paxinos and Watson, 1998).

#### Classical trace eyeblink conditioning and signal processing

Each rat was conditioned using a white noise (75 dB, 200 ms, Fig. 1B) as a conditioned stimulus (CS) and a 100-Hz burst of 0.5 ms bipolar pulses of periorbital shocks (100 ms) as an unconditioned stimulus (US). The amplitude of the US was adjusted to individually for each rat to elicit a full blinking of the eye. Each trial started with the 200-ms CS presentation, followed by a 500-ms stimulus-free trace period, and then a 100-ms shock-US. Each rat performed 8 sessions and each session consisted of 60 trials.

### Signal pre-processing

#### Artefact removal

We analyzed a 500-ms sweeps of LFP signal starting from the CS onset. Trials with large LFP fluctuations due to movement or device-related artefacts were excluded based on simple thresholding with a 1500 µV cutoff. This procedure resulted in a rejection rate of 0.6%. At the time of CS onset (0ms) and offset (200ms), we observed very short-lived (<3ms) power increase in high frequency ranges (>100Hz) as potentially caused by stimulus driven artefacts. To remove the spike artefact, signals from -2.5ms to 1.5ms around CS onset/offset were replaced by preceding spontaneous signal in each individual trial. In the analyses of comparing perceived vs. unperceived trials and comparing sessions of highest vs. lowest hit rate, we further excluded trials with noisy EMG trace, and balanced the number of trials between the two conditions. In this process, we randomly selected trials to match the minimum number of trials within a session.

#### LFP re-referencing and filtering

We applied CSD re-referencing to raw LFP by subtracting the signal by the signal average of two adjacent channels to remove any possible common noise that all the electrodes would have picked up. The results, however, did not differ from using raw signal (no re-referencing). Here, we report results using re-referenced signal. We used finite impulse response (FIR) filter composed of high-pass stop bands of 0.3 and low-pass stop bands of 1.4 to filter frequencies with log scaled increment to compute amplitude, phase-locking to stimulus and inter-areal 1:1 synchrony. We filtered frequencies from 3Hz to 45Hz with a step of 1Hz to compute CFCs. The filtered signal was Hilbert transformed to obtain the imaginary part of the complex vector (Palva et al., 2005).

### Analysis of conditioned eyeblink responses

Eyeblinks were detected from the EMG signal offline to determine the percentage of conditioned responses (same as hit rate, HR). Briefly, the EMG signal during the last 200 ms of the trace period was analyzed (Fig. 1B, time window is shaded in gray color). The same analysis method was used in detecting blinking as previously reported (Nokia et al., 2017). The HR was obtained by counting the number of trials during which the rat blinked divided by the total number of trials in a session.

### Time-frequency representation of amplitude and stimulus locking across trials

We used the filtered signal to measure amplitude and compute stimulus locking (phase locking value) to CS onset across trials, which were represented in time frequency matrix. To obtain relative change in amplitude and stimulus locking to baseline activity, we subtracted the amplitude and stimulus locking by that of averaged value during baseline period (from 600 ms to100 ms prior to CS onset).

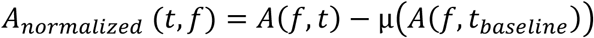

where time series of LFP amplitude is expressed as *A*(*f, t*) at a given frequency of *f* and time *t*, mean, µ(*A*(*f, t*_*baseline*_)) of the baseline period.

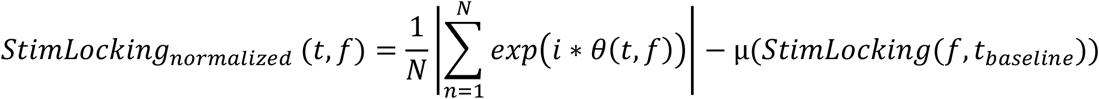

where *i* is the imaginary unit, and phase time series *θ*(*t, f*) across N number of trials at a given frequency *f*. Then, *StimLocking*_*normalized*_ (*t, f*) was derived by subtracting it using the baseline average µ(*StimLocking*(*f, t*_*baseline*_)).

#### 1:1 phase synchrony and phase-amplitude coupling

1:1 phase synchrony between fissure and hilus was derived by using phase time series *θ*(*t, f*)_*fis*_, and *θ*(*t, f*)_*hil*_, respectively, in each given frequency *f*. During this process, we normalized the PLV by dividing the value by the average of 100 surrogate PLV obtained by shuffling trials to control contributions of artificial PLV driven by stimulus-locked component.

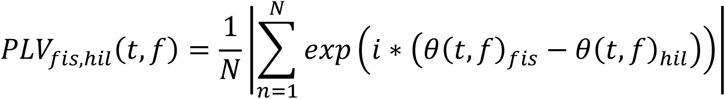

We further quantified *n:m* (frequency ratio) phase-amplitude coupling (PAC), one way to compute cross frequency coupling using phase locking value between pairs of recorded regions (within fissure, *ff*; fissure-hilus, *fh*; hilus-fissure, *hf*; within hilus, *hh*) and between the phase of the slow oscillation (LF) and the phase of the amplitude envelope, *θ*^*E*^(*t, HF*) of the HF filtered at LF (Vanhatalo et al., 2004)at each of the laminar pair. We used integers n = 1, and m = 2 to 9 with a step of 1 as the frequency ratios. (Palva et al., 2005; Siebenhühner et al., 2016). The same normalization was applied as it was done in 1:1 inter areal synchrony analysis using surrogates obtained by trial shuffling.

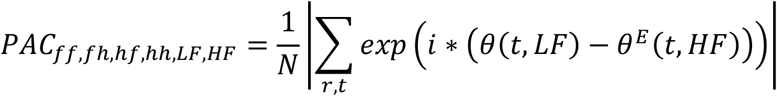

where *i, r, t*, imaginary unit, number of trials, and number of samples t within a window, respectively. PAC was computed for pre- and post-CS period, and the difference between the two was used to evaluate synchronization. We evaluated the results for separate analysis time windows; 0-100 ms, 100–300 ms, 300–500 ms after CS onset.

### Phase transfer entropy

As described in Lobier et al., (2014), phase TE was derived by using the instantaneous phase time-series of the signal X _hilus_(*t*) and Y _fissure_(*t*) expressed as *θ* _hilus_(*t*) and *θ* _fissure_(*t*) after complex filtering of the signal using Hilbert transform (Le Van Quyen et al., 2001) for a given frequency band. In the analysis using empirical data, we used 500ms of post-stimulus period and delay lag (*δ*) was defined as proportional to the number of cycles in the signal for a given frequency.

Phase TE _hil→fis_ = H (θ_fis_(t), θ_fis_(t’)) + H (θ_fis_(t’), θ_hil_(t’))-H (θ_fis_(t’))- H (θ_fis_(t), θ_fis_(t’), θ_hil_(t’))where *θ*_*hil*_(*t’*) and *θ*_*fis*_(*t’*) denote past states at time point *t’ = t-* δ, thus, *θ*_*hil*_(*t’*) = *θ*_*hil*_ (*t-* δ) and *θ*_*fis*_(*t’*) = *θ*_*fis*_ (*t-* δ). The marginal entropy and joint entropy are defined as below: H (θ_fis_(t), θ_fis_(t’)) =−Σp(θ_fis_(t), θ_fis_(t’)) x log p(θ_fis_(t), θ_fis_(t’))

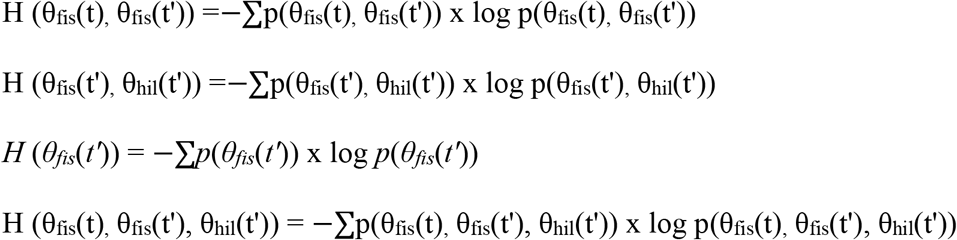

In our analysis, we used pairs of phase data points for each computation where we used the binning method described in Lobier et al., (2014) with the bin width defined following Scott’s choice (Scott, 1992). To create bias-free measure, we computed differential TE (dTE) derived by pTE _hil→fis_ - pTE _fis→hil_and used this measure for further analysis. We next obtained surrogate data sets (n=1000) by trial shuffling and compared with the empirical data to test the statistical significance against a null hypothesis of no information transfer between hilus and fissure (Monte Carlo *p*<0.05).

### Statistical analysis

To obtain whether neural activity (amplitude, stimulus locking, and 1:1 inter-areal synchrony) was significantly increased or decreased in response to CS, we used a coin flip permutation procedure. We compared the normalized data that was baseline subtracted with a distribution derived by that of permuted data (20,000) using random flips with probability of 0.5. Then, Monte-Carlo *p*<0.025 (two-tailed), was obtained, where further corrected for multiple comparisons using the Hochberg and Benjamini procedure (Hochberg and Benjamini, 1990) in each analysis time window.

To assess significant difference in oscillation amplitudes, stimulus locking across trials, and 1:1 inter-areal synchrony between sessions with the highest and lowest hit rate, and between perceived and unperceived trials, we first obtained baseline subtracted time-frequency representation of neural activity. Using this data, we ran paired t-test between sessions with highest and lowest hit rate across subjects and obtained empirical *t* value corresponding to a *p* value with a first level t-threshold (*p*<0.025, two-tailed) for each time *t* and frequency *f*, clustered together based on temporal adjacency at each frequency. While retaining the maximum t-sum observed value, we created 1000 surrogate sessions by randomly assigning sessions with highest and lowest hit rate and repeated clustering procedure on the surrogate dataset using cluster –based permutation *t* test (Maris and Oostenveld, 2007).

Thereafter we derived Monte-Carlo probability distribution at *p*<0.05 by comparing the observed maximum t-sum and maximum t-sum distribution obtained under the null hypothesis. The same clustering permutation test was performed for testing significant difference between perceived (CR) and unperceived (no-CR) trials as well as trials with largest and smallest HR.

The presence of significant PAC in the post-stimulus latencies was assessed using one sample t-test across all rats and all sessions in each low frequency, *n:m* ratio, analysis time window, and laminar pair. Multiple comparisons were then corrected using Hochberg and Benjamini (1990) method.

To evaluate significance of PAC difference between sessions showing the highest and lowest hit rate, or between perceived and unperceived trials, we performed paired t-test (*p*<0.01) for *n:m* pairs showing significant modulation of PAC during post-stimulus period.

## References

Akam T, Oren I, Mantoan L, Ferenczi E, Kullmann DM. 2012. Oscillatory dynamics in the hippocampus support dentate gyrus–CA3 coupling. Nat Neurosci 15:763–768.

Andersen P, Morris R, Amaral D, Bliss T, O’Keefe J. 2006. The hippocampus book. Oxford university press.

Bahramisharif A, Jensen O, Jacobs J, Lisman J. 2018. Serial representation of items during working memory maintenance at letter-selective cortical sites. PLoS Biol 16:e2003805.

Belluscio MA, Mizuseki K, Schmidt R, Kempter R, Buzsáki G. 2012. Cross-frequency phase–phase coupling between theta and gamma oscillations in the hippocampus. J Neurosci 32:423–435.

Berry SD, Thompson RF. 1978. Prediction of learning rate from the hippocampal electroencephalogram. Science (80-) 200:1298–1300.

Buzsáki G. 2002. Theta oscillations in the hippocampus. Neuron 33:325–340.

Buzsáki G. 1989. Two-stage model of memory trace formation: a role for “noisy” brain states. Neuroscience 31:551–570.

Buzsáki G, Chrobak JJ. 1995. Temporal structure in spatially organized neuronal ensembles: a role for interneuronal networks. Curr Opin Neurobiol 5:504–510.

Buzsáki G, Moser EI. 2013. Memory, navigation and theta rhythm in the hippocampal-entorhinal system. Nat Neurosci 16:130–138.

Canolty RT, Knight RT. 2010. The functional role of cross-frequency coupling. Trends Cogn Sci 14:506–515.

Caplan JB, Madsen JR, Schulze-Bonhage A, Aschenbrenner-Scheibe R, Newman EL, Kahana MJ. 2003. Human θ oscillations related to sensorimotor integration and spatial learning. J Neurosci 23:4726–4736.

Cason H. 1922. The conditioned eyelid reaction. J Exp Psychol 5:153.

Cheng DT, Disterhoft JF, Power JM, Ellis DA, Desmond JE. 2008. Neural substrates underlying human delay and trace eyeblink conditioning. Proc Natl Acad Sci 105:8108–8113.

Chrobak JJ, Buzsáki G. 1996. High-frequency oscillations in the output networks of the hippocampal–entorhinal axis of the freely behaving rat. J Neurosci 16:3056–3066.

Clark RE, Manns JR, Squire LR. 2002. Classical conditioning, awareness, and brain systems. Trends Cogn Sci 6:524–531.

Clark RE, Squire LR. 1998. Classical conditioning and brain systems: The role of awareness. Science (80-). doi:10.1126/science.280.5360.77

Colgin LL. 2016. Rhythms of the hippocampal network. Nat Rev Neurosci 17:239–249.

Colgin LL. 2013. Mechanisms and functions of theta rhythms. Annu Rev Neurosci 36:295–312.

Colgin LL. 2011. Oscillations and hippocampal–prefrontal synchrony. Curr Opin Neurobiol 21:467– 474.

Colgin LL, Denninger T, Fyhn M, Hafting T, Bonnevie T, Jensen O, Moser M-B, Moser EI. 2009. Frequency of gamma oscillations routes flow of information in the hippocampus. Nature 462:353–357.

Das A, Menon V. 2021. Asymmetric frequency-specific feedforward and feedback information flow between hippocampus and prefrontal cortex during verbal memory encoding and recall. J Neurosci 41:8427–8440.

DeCoteau WE, Thorn C, Gibson DJ, Courtemanche R, Mitra P, Kubota Y, Graybiel AM. 2007. Learning-related coordination of striatal and hippocampal theta rhythms during acquisition of a procedural maze task. Proc Natl Acad Sci 104:5644–5649.

Dehaene S, Changeux J-P. 2011. Experimental and theoretical approaches to conscious processing. Neuron 70:200–227.

Del Cul A, Baillet S, Dehaene S. 2007. Brain dynamics underlying the nonlinear threshold for access to consciousness. PLoS Biol 5:e260.

Fell J, Axmacher N. 2011. The role of phase synchronization in memory processes. Nat Rev Neurosci 12:105–118.

Fernández-Ruiz A, Oliva A, Nagy GA, Maurer AP, Berényi A, Buzsáki G. 2017. Entorhinal-CA3 dual-input control of spike timing in the hippocampus by theta-gamma coupling. Neuron 93:1213–1226.

Gloveli T, Schmitz D, Empson RM, Heinemann U. 1997. Frequency-dependent information flow from the entorhinal cortex to the hippocampus. J Neurophysiol 78:3444–3449.

Griffiths BJ, Martín -Buro Mc, Staresina BP, Hanslmayr S. 2021. Disentangling neocortical alpha/beta and hippocampal theta/gamma oscillations in human episodic memory formation. Neuroimage 242:118454.

Griffiths BJ, Mayhew SD, Mullinger KJ, Jorge J, Charest I, Wimber M, Hanslmayr S. 2019. Alpha/beta power decreases track the fidelity of stimulus-specific information. Elife 8:e49562.

Hirvonen J, Palva S. 2016. Cortical localization of phase and amplitude dynamics predicting access to somatosensory awareness. Hum Brain Mapp 37:311–326.

Hochberg Y, Benjamini Y. 1990. More powerful procedures for multiple significance testing. Stat Med 9:811–818. doi:10.1002/sim.4780090710

Hyafil A, Giraud A-L, Fontolan L, Gutkin B. 2015. Neural cross-frequency coupling: connecting architectures, mechanisms, and functions. Trends Neurosci 38:725–740.

Igarashi KM, Lu L, Colgin LL, Moser M-B, Moser EI. 2014. Coordination of entorhinal– hippocampal ensemble activity during associative learning. Nature 510:143–147.

Jensen O, Bonnefond M, Marshall TR, Tiesinga P. 2015. Oscillatory mechanisms of feedforward and feedback visual processing. Trends Neurosci 38:192–194.

Julku H, Rouhinen S, Huttunen HJ, Lindberg L, Liinamaa J, Saarela V, Karvonen E, Booms S, Mäkelä JP, Uusitalo H. 2021. Reduced evoked activity and cortical oscillations are correlated with anisometric amblyopia and impairment of visual acuity. Sci Rep 11:1–15.

Jutras MJ, Fries P, Buffalo EA. 2013. Oscillatory activity in the monkey hippocampus during visual exploration and memory formation. Proc Natl Acad Sci 110:13144–13149.

Jutras MJ, Fries P, Buffalo EA. 2009. Gamma-band synchronization in the macaque hippocampus and memory formation. J Neurosci 29:12521–12531.

Kay LM. 2014. Circuit oscillations in odor perception and memory. Prog Brain Res 208:223–251.

Kreiman G, Fried I, Koch C. 2002. Single-neuron correlates of subjective vision in the human medial temporal lobe. Proc Natl Acad Sci U S A 99:8378–8383. doi:10.1073/pnas.072194099

Lamme VAF, Roelfsema PR. 2000. The distinct modes of vision offered by feedforward and recurrent processing. Trends Neurosci 23:571–579.

Lamy D, Salti M, Bar-Haim Y. 2009. Neural correlates of subjective awareness and unconscious processing: an ERP study. J Cogn Neurosci 21:1435–1446.

Le Van Quyen M, Foucher J, Lachaux J-P, Rodriguez E, Lutz A, Martinerie J, Varela FJ. 2001. Comparison of Hilbert transform and wavelet methods for the analysis of neuronal synchrony. J Neurosci Methods 111:83–98.

Lee ACH, Yeung L-K, Barense MD. 2012. The hippocampus and visual perception. Front Hum Neurosci 6:91.

Lisman JE. 1999. Relating hippocampal circuitry to function: recall of memory sequences by reciprocal dentate–CA3 interactions. Neuron 22:233–242.

Lisman JE, Jensen O. 2013. The theta-gamma neural code. Neuron 77:1002–1016.

Lobier M, Siebenhühner F, Palva S, Palva JM. 2014. Phase transfer entropy: a novel phase-based measure for directed connectivity in networks coupled by oscillatory interactions. Neuroimage 85:853–872.

Mansvelder HD, Verhoog MB, Goriounova NA. 2019. Synaptic plasticity in human cortical circuits: cellular mechanisms of learning and memory in the human brain? Curr Opin Neurobiol 54:186– 193.

Maris E, Oostenveld R. 2007. Nonparametric statistical testing of EEG- and MEG-data. J Neurosci Methods 164:177–190. doi:10.1016/j.jneumeth.2007.03.024

McEchron MD, Disterhoft JF. 1999. Hippocampal encoding of non-spatial trace conditioning. Hippocampus 9:385–396.

Michalareas G, Vezoli J, Van Pelt S, Schoffelen J-M, Kennedy H, Fries P. 2016. Alpha-beta and gamma rhythms subserve feedback and feedforward influences among human visual cortical areas. Neuron 89:384–397.

Morgan LK, MacEvoy SP, Aguirre GK, Epstein RA. 2011. Distances between real-world locations are represented in the human hippocampus. J Neurosci 31:1238–1245.

Nokia MS, Gureviciene I, Waselius T, Tanila H, Penttonen M. 2017. Hippocampal electrical stimulation disrupts associative learning when targeted at dentate spikes. J Physiol 595:4961– 4971.

Nokia MS, Penttonen M, Korhonen T, Wikgren J. 2009. Hippocampal theta-band activity and trace eyeblink conditioning in rabbits. Behav Neurosci 123:631.

Nokia MS, Sisti HM, Choksi MR, Shors TJ. 2012. Learning to learn: theta oscillations predict new learning, which enhances related learning and neurogenesis. PLoS One 7:e31375.

Nokia MS, Waselius T, Mikkonen JE, Wikgren J, Penttonen M. 2015. Phase matters: responding to and learning about peripheral stimuli depends on hippocampal θ phase at stimulus onset. Learn Mem 22:307–317.

O’Keefe J, Dostrovsky J. 1971. The hippocampus as a spatial map: preliminary evidence from unit activity in the freely-moving rat. Brain Res.

O’keefe J, Nadel L. 1978. The hippocampus as a cognitive map. Oxford university press.

Palva JM, Palva S, Kaila K. 2005. Phase synchrony among neuronal oscillations in the human cortex. J Neurosci 25:3962–3972.

Palva S, Kulashekhar S, Hämäläinen M, Palva JM. 2011. Localization of cortical phase and amplitude dynamics during visual working memory encoding and retention. J Neurosci 31:5013–5025.

Paxinos G, Watson C. 1998. A stereotaxic atlas of the rat brain. New York Acad.

Rangel LM, Rueckemann JW, Riviere PD, Keefe KR, Porter BS, Heimbuch IS, Budlong CH, Eichenbaum H. 2016. Rhythmic coordination of hippocampal neurons during associative memory processing. Elife 5:e09849.

Reber TP, Faber J, Niediek J, Boström J, Elger CE, Mormann F. 2017. Single-neuron correlates of conscious perception in the human medial temporal lobe. Curr Biol 27:2991–2998.

Richter CG, Thompson WH, Bosman CA, Fries P. 2017. Top-down beta enhances bottom-up gamma. J Neurosci 37:6698–6711.

Scharfman HE. 2016. The enigmatic mossy cell of the dentate gyrus. Nat Rev Neurosci 17:562–575.

Scheffer-Teixeira R, Belchior H, Caixeta F V, Souza BC, Ribeiro S, Tort ABL. 2012. Theta phase modulates multiple layer-specific oscillations in the CA1 region. Cereb cortex 22:2404–2414.

Schneiderman N, Fuentes I, Gormezano I. 1962. Acquisition and extinction of the classically conditioned eyelid response in the albino rabbit. Science (80-) 136:650–652.

Schroeder CE, Lakatos P. 2009. Low-frequency neuronal oscillations as instruments of sensory selection. Trends Neurosci 32:9–18.

Scott DW. 2015. Multivariate density estimation: theory, practice, and visualization. John Wiley & Sons.

Scoville WB, Milner B. 1957. Loss of recent memory after bilateral hippocampal lesions. J Neurol Neurosurg Psychiatry 20:11.

Seager MA, Johnson LD, Chabot ES, Asaka Y, Berry SD. 2002. Oscillatory brain states and learning: Impact of hippocampal theta-contingent training. Proc Natl Acad Sci 99:1616–1620.

Sergent C, Baillet S, Dehaene S. 2005. Timing of the brain events underlying access to consciousness during the attentional blink. Nat Neurosci 8:1391–1400. doi:10.1038/nn1549

Siebenhühner F, Wang SH, Palva JM, Palva S. 2016. Cross-frequency synchronization connects networks of fast and slow oscillations during visual working memory maintenance. Elife 5:e13451.

Singer W. 1995. Development and plasticity of cortical processing architectures. Science (80-) 270:758–764.

Squire LR. 1992. Memory and the hippocampus: A synthesis from findings with rats, monkeys, and humans. Psychol Rev. doi:10.1037/0033-295X.99.2.195

Takehara-Nishiuchi K, Maal-Bared G, Morrissey M. 2012. Increased entorhinal–prefrontal theta synchronization parallels decreased entorhinal–hippocampal theta synchronization during learning and consolidation of associative memory. Front Behav Neurosci 5:90.

Teyler TJ, Rudy JW. 2007. The hippocampal indexing theory and episodic memory: updating the index. Hippocampus 17:1158–1169.

Tort ABL, Komorowski R, Eichenbaum H, Kopell N. 2010. Measuring phase-amplitude coupling between neuronal oscillations of different frequencies. J Neurophysiol 104:1195–1210.

Tort ABL, Komorowski RW, Manns JR, Kopell NJ, Eichenbaum H. 2009. Theta–gamma coupling increases during the learning of item–context associations. Proc Natl Acad Sci 106:20942– 20947.

Tort ABL, Kramer MA, Thorn C, Gibson DJ, Kubota Y, Graybiel AM, Kopell NJ. 2008. Dynamic cross-frequency couplings of local field potential oscillations in rat striatum and hippocampus during performance of a T-maze task. Proc Natl Acad Sci 105:20517–20522.

Treder M, Charest I, Michelmann S, Martín-Buro MC, Roux F, Carceller-Benito F, Ugalde-Canitrot A, Rollings D, Sawlani V, Chelvarajah R. 2021. The hippocampus as the switchboard between perception and memory. bioRxiv 2005–2020.

Tseng W, Guan R, Disterhoft JF, Weiss C. 2004. Trace eyeblink conditioning is hippocampally dependent in mice. Hippocampus 14:58–65.

Uhlhaas PJ, Roux F, Rodriguez E, Rotarska-Jagiela A, Singer W. 2010. Neural synchrony and the development of cortical networks. Trends Cogn Sci 14:72–80.

Urgolites ZJ, Smith CN, Squire LR. 2018. Eye movements support the link between conscious memory and medial temporal lobe function. Proc Natl Acad Sci 115:7599–7604.

Urgolites ZJ, Wixted JT, Goldinger SD, Papesh MH, Treiman DM, Squire LR, Steinmetz PN. 2020. Spiking activity in the human hippocampus prior to encoding predicts subsequent memory. Proc Natl Acad Sci 117:13767–13770.

Vanhatalo S, Palva JM, Holmes MD, Miller JW, Voipio J, Kaila K. 2004. Infraslow oscillations modulate excitability and interictal epileptic activity in the human cortex during sleep. Proc Natl Acad Sci 101:5053–5057.

Voigt MB, Yusuf PA, Kral A. 2018. Intracortical microstimulation modulates cortical induced responses. J Neurosci 38:7774–7786.

Wallenstein G V, Hasselmo ME, Eichenbaum H. 1998. The hippocampus as an associator of discontiguous events. Trends Neurosci 21:317–323.

Wang X-J, Buzsáki G. 1996. Gamma oscillation by synaptic inhibition in a hippocampal interneuronal network model. J Neurosci 16:6402–6413.

Weiss C, Kronforst-Collins MA, Disterhoft JF. 1996. Activity of hippocampal pyramidal neurons during trace eyeblink conditioning. Hippocampus 6:192–209.

Whittington MA, Traub RD, Jefferys JGR. 1995. Synchronized oscillations in interneuron networks driven by metabotropic glutamate receptor activation. Nature 373:612–615.

Wikgren J, Nokia MS, Penttonen M. 2010. Hippocampo–cerebellar theta band phase synchrony in rabbits. Neuroscience 165:1538–1545.

Winson J. 1978. Loss of hippocampal theta rhythm results in spatial memory deficit in the rat. Science (80-) 201:160–163.

Zeidman P, Mullally SL, Maguire EA. 2015. Constructing, perceiving, and maintaining scenes: hippocampal activity and connectivity. Cereb Cortex 25:3836–3855.

Zheng C, Bieri KW, Hwaun E, Colgin LL. 2016. Fast gamma rhythms in the hippocampus promote encoding of novel object–place pairings. Eneuro 3.

